# Deep and fast label-free Dynamic Organellar Mapping

**DOI:** 10.1101/2021.11.09.467934

**Authors:** Julia P. Schessner, Vincent Albrecht, Alexandra K. Davies, Pavel Sinitcyn, Georg H.H. Borner

## Abstract

The Dynamic Organellar Maps (DOMs) approach combines cell fractionation and shotgun-proteomics for global profiling analysis of protein subcellular localization. Here, we have drastically enhanced the performance of DOMs through data-independent acquisition (DIA) mass spectrometry (MS). DIA-DOMs achieve twice the depth of our previous workflow in the same MS runtime, and substantially improve profiling precision and reproducibility. We leveraged this gain to establish flexible map formats scaling from rapid analyses to ultra-deep coverage. Our fastest format takes only ∼2.5h/map and enables high-throughput experimental designs. Furthermore, we introduce DOM-QC, an open-source software tool for in-depth standardized analysis of DOMs and other profiling data. We then applied DIA-DOMs to capture subcellular localization changes in response to starvation and disruption of lysosomal pH in HeLa cells, which revealed a subset of Golgi proteins that cycle through endosomes. DIA-DOMs offer a superior workflow for label-free spatial proteomics as a systematic phenotype discovery tool.

## Introduction

The compartments of eukaryotic cells organize the proteome into dynamic reaction spaces that control protein activity. The large number of diseases caused by disrupted protein transport demonstrates that protein localization must be tightly regulated to ensure correct protein function [1,2]. Our understanding of cellular homeostasis thus requires a comprehensive view of protein localizations and movements within the cell, collectively termed the ‘spatial proteome’ [3–10].

Our lab previously developed the Dynamic Organellar Maps (DOMs) approach for systems-level capture of protein subcellular localization [11]. Briefly, cells are lysed mechanically and the released organelles are partially separated by differential centrifugation [11,12]. Pelleted proteins are quantified across the fractions by mass spectrometry (MS). The obtained abundance profiles are characteristic of the harboring organelles and are used to predict protein localization by supervised machine-learning; the result is an organellar ‘map of the cell’ [11]. Since DOMs are highly reproducible, they allow capture of induced protein localization changes, and have thus driven phenotype discovery in diverse biological contexts. For example, DOMs have been applied to reveal the molecular pathomechanisms of AP-4 deficiency syndrome, a severe neurological disorder [13,14]; to characterize the function of a lysosomal retrieval pathway [15]; to quantify translocation events triggered during EGF signalling [11]; to identify the target of drugs selected from a phenotypic screen [16]; and to uncover how HIV infection alters the composition of extracellular vesicles [17].

The original DOMs method relied on SILAC (stable isotope labelling by amino acids in cell culture) [18] for accurate protein quantification across subcellular fractions [11]. To extend the method beyond cell lines amenable to SILAC, we implemented different quantification strategies [4,19], including label-free quantification (LFQ) [20] and the peptide-labelling methods TMT [21] and EASI-tag [22]. SILAC-based maps yield the most precise profiles, but offer limited depth due to increased MS1 spectral complexity. LFQ maps achieve greater depth but suffer from lower precision, while TMT and EASI-tag maps have intermediate quality [4].

All aforementioned DOMs workflows are based on data-dependent acquisition (DDA) of MS data [23,24]. In each DDA cycle, a fixed number of the most abundant peptides from the MS1 scan are individually fragmented for identification. The MS1 intensities are subsequently used for quantification. While this approach prioritizes precursors that are most likely to generate high-quality MS2 spectra, it adds a stochastic element to precursor selection, which leads to inconsistent protein identifications across samples. In the context of DOMs, missing values severely limit depth of analysis, since profiling requires quantification of the same protein in the majority of measured subcellular fractions and replicates. To alleviate this problem, we previously fractionated peptide samples prior to MS analysis [11,25]. This resulted in improved map depth, but tripled MS time requirements.

Owing to recent advances in MS instrumentation and data analysis software, data-independent acquisition (DIA) is increasingly replacing DDA approaches [26]. During DIA, the entire peptide mass range of the MS1 scan is partitioned into windows. Co-eluting peptides within each window are fragmented together, resulting in complex MS2 spectra. Their deconvolution requires an auxiliary library with reference peptide spectra, which can either be generated from *in silico* predictions, from additionally measured DDA spectra, or directly from the DIA data [27]. The DIA approach is technically and computationally challenging, but conceptually allows the identification of all peptides present in a sample. Moreover, unlike in DDA, both the MS1 precursor and the MS2 fragment ions can be used for quantification, which increases precision [28]. DIA is hence becoming the strategy of choice for extensive profiling-based approaches such as SEC-MS [29] and has recently been applied in high-throughput subcellular phosphoproteomics [9].

Here, we harness the power of DIA for generating label-free DOMs. We show that proteomic depth, precision and reproducibility of DIA-DOMs increase dramatically relative to DDA-DOMs. For this purpose, we introduce the software tool DOM-QC, which enables rapid standardized analysis of DOMs and other types of profiling data. We provide optimized DIA-DOMs formats with short MS run times suitable for high-throughput experiments and with longer MS runtimes for maximum coverage. Finally, we investigate subcellular rearrangements upon starvation and inhibition of lysosomal acidification in HeLa cells, to demonstrate the power of DIA-DOMs for phenotype discovery.

## Results

### DOM-QC, a new software tool for in-depth analysis and benchmarking of profiling data

To facilitate the establishment and optimization of a new DIA-DOMs workflow, we first implemented DOM-QC, a software tool for the exploration and benchmarking of profiling data. DOM-QC provides multiple metrics and interactive info-plots via a graphical user interface (https://domqc.bornerlab.org). As input data, the tool handles raw output files from MaxQuant [30] and Spectronaut [31], as well as any pre-processed profiling data. Multiple maps can be uploaded together and directly compared. Proteomic depth is assessed before and after filtering for usable profiles, per map, and across replicates. Principal component analysis (PCA) plots provide a visual overview of map topology. Three metrics are calculated to gauge map quality: 1. Profile scatter of proteins that are part of the same complex. This reflects within-map profiling precision, based on the assumption that tightly bound proteins co-fractionate with near-identical profiles. 2. Profile scatter of individual proteins across map replicates, which reflects inter-map reproducibility. 3. Organellar prediction performance (supervised machine learning by support vector machines (SVMs)), judged by the harmonic mean of recall and precision of pre-defined marker proteins (F1 score).

As a benchmark sample set for the optimization of DIA-DOMs, we prepared three independent subcellular fractionations from HeLa cells, each with six fractions, label-free, as described [19]. We then tested a variety of liquid chromatography-mass spectrometry (LC-MS) setups and data acquisition strategies for mass spectrometric analysis (see Fig. 1 for overview of the experimental design and Supplementary Fig. 1 for DIA method optimization). Data were processed with MaxQuant 2.0, which features the new MaxDIA algorithm [32].

**Fig. 1.**
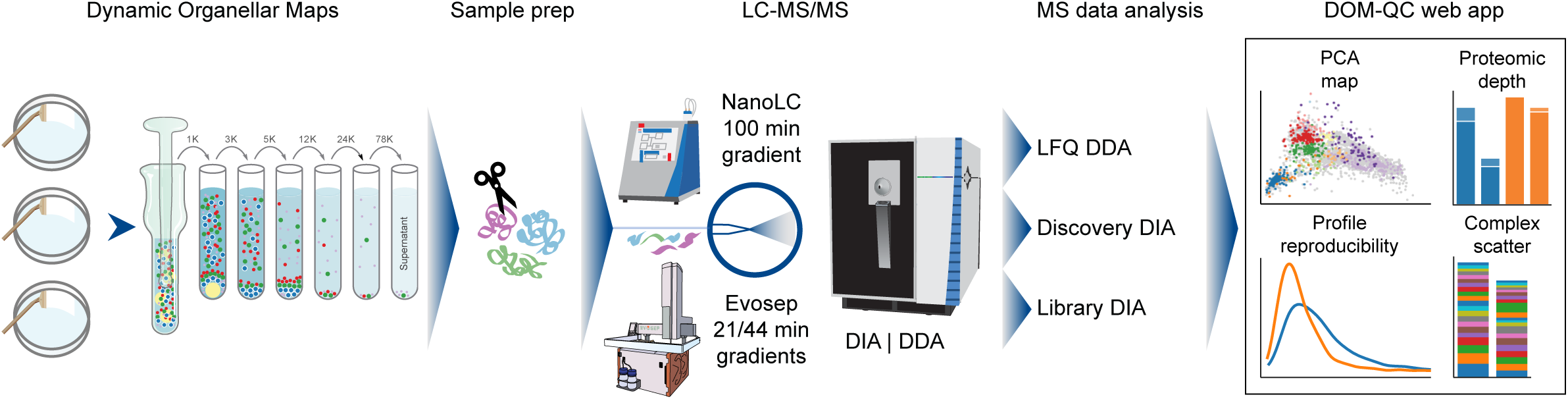
Overview of the Dynamic Organellar Maps workflow, DIA optimization strategy, and evaluation with DOM-QC. HeLa cells were fractionated by differential centrifugation (x g indicated), to generate triplicate reference organellar maps. Following tryptic digest, samples were analyzed by LC-MS, using either a Thermo EASY-nLC 1200 HPLC, or a high-throughput Evosep One HPLC, coupled to a Thermo Exploris 480 orbitrap mass spectrometer. Data were acquired in DDA or DIA mode, and DIA processing was performed with library or discovery DIA. For the performance evaluation, data were analyzed with the new web app DOM-QC (https://domqc.bornerlab.org), which provides multiple info graphics and metrics for assessing map topology, proteomic depth, map resolution and reproducibility.

### DIA maps outperform DDA maps across all metrics

To compare DIA-DOMs to our previous label-free DDA-DOMs workflow, we measured our benchmark samples with DIA and DDA, on an Orbitrap Exploris mass spectrometer fitted with a nanoLC (100 min gradient). DIA data were processed either with a custom spectral library (ca. unique 159,000 peptide sequences, acquired with DDA), or with an *in-silico* spectral library predicted by DeepMassPrism [33] (MaxQuant’s ‘discovery DIA’ mode [32]). PCA plots of the DDA-, library DIA- and discovery DIA-based maps looked topologically similar (Fig. 2A), but enhanced tightness of organellar clusters was apparent in the DIA maps. As expected, the unfiltered proteome depth was only slightly increased by DIA (7,067 protein groups (PGs) with DDA vs. 7,371 PGs with discovery DIA and 7,833 PGs with library DIA). In contrast, the number of proteins profiled across all three map replicates dramatically increased from 2,768 PGs with DDA, to 5,241 PGs with discovery DIA (+89%), and to 6,719 PGs with library DIA (+143%; Fig. 2B). This performance leap is explained by the much more consistent identification of proteins across samples, reflected by a rise in data completeness from 69% with DDA to 90% with discovery DIA, and to 97% with library DIA (Supplementary Fig. 2A). The proteins profiled in the DIA datasets mostly overlapped and contained almost all proteins quantified by DDA (Fig. 2B). Next, we assessed SVM-based organellar classification, analyzing a set of 844 established organellar marker proteins common to all three maps. DIA maps moderately but clearly outperformed DDA maps. Overall recall increased from 93% (DDA) to 95% (DIA) and the average F1 score increased from 0.87 to 0.9 (Fig. 2C). Importantly, the F1 scores for the highly dynamic membrane compartments endoplasmic reticulum (ER), plasma membrane, lysosomes and endosomes improved substantially with DIA (Supplementary Fig. 2B). The combination of increased depth and better SVM performance also resulted in a greatly increased number of high-confidence localization predictions, up from 998 (non-marker proteins) with DDA, to 1,649 with discovery DIA (+65%), and to 2,112 with library DIA (+111%) (Fig. 2D). In addition, DIA maps also provided many more medium-confidence predictions (Fig. 2D). For all three datasets, the concordance with our previously published predictions based on SILAC DOMs [11] was 98-100% in the high-confidence category (Supplementary Fig. 2C). Finally, we evaluated profile quantification precision and reproducibility, which are key for comparative spatial proteomics [4]. Within-map profiling precision was markedly improved with DIA (complex scatter reduced by 13% with discovery DIA and by 19% with library DIA; Fig. 2E), and inter-map profile reproducibility was greatly enhanced (profile scatter reduced by 24% with discovery DIA and by 35% with library DIA; Fig. 2F). Taken together, these data demonstrate that DIA-DOMs strongly outperform our previously established label free DDA-DOMs with regards to depth, organellar resolution, precision and reproducibility. Remarkably, this drastic improvement is already achieved with discovery DIA using an *in silico* predicted spectral library, and can be boosted even further by using a measured peptide library.

**Fig. 2.**
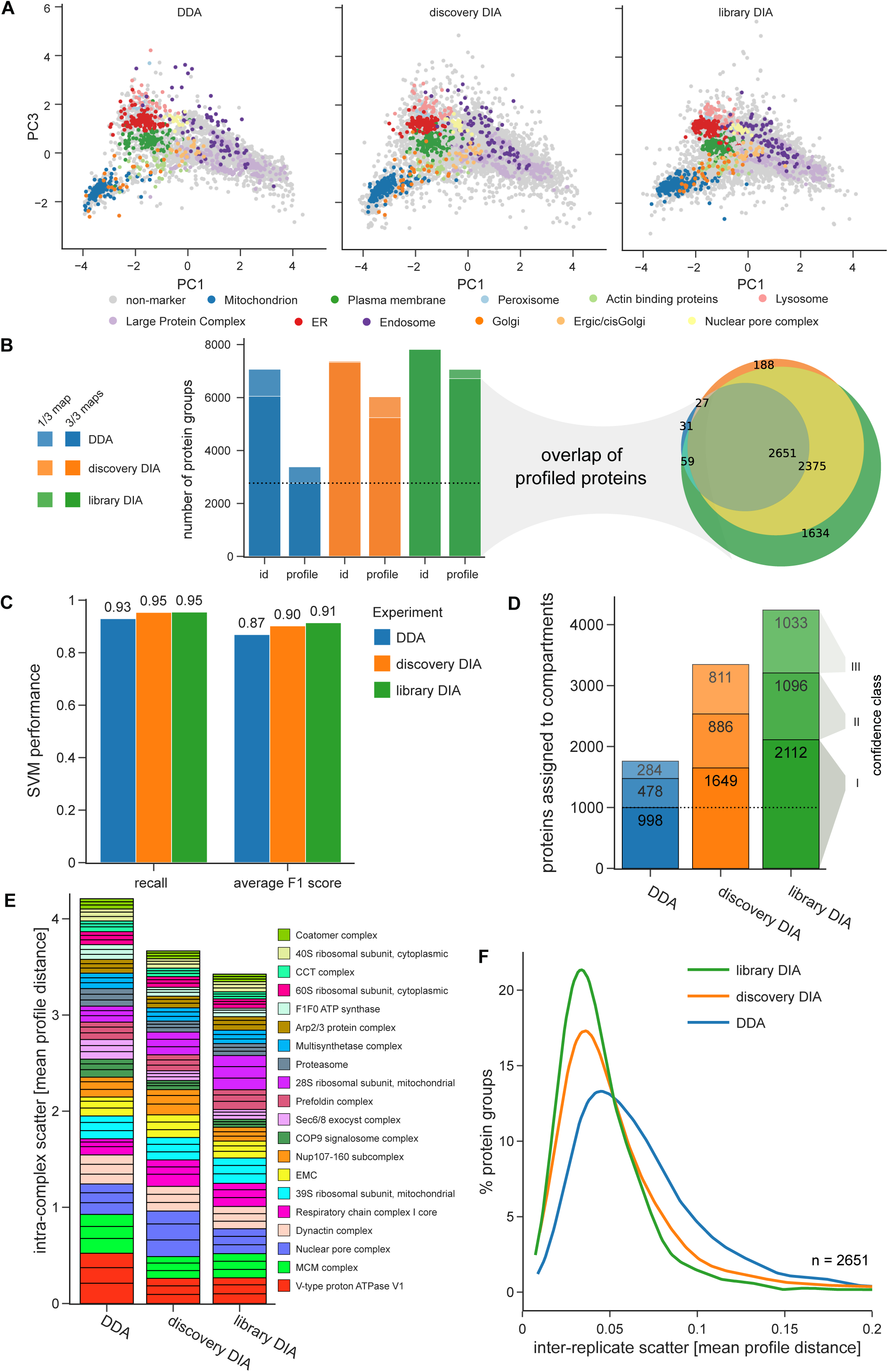
Comparison of DDA, discovery DIA and library DIA based maps. All maps were acquired with 100 min LC gradients. **A)** Topology of organellar maps in PCA space. Coloured dots correspond to organellar marker proteins. A single PCA was performed across all three experiments. PCs 1 and 3 provide the best visual separation of clusters. **B)** Left panel: Number of proteins identified or profiled in at least 1 or in all 3 out of 3 replicate maps. Right panel: Overlap of profiled proteins between acquisition modes. **C)** Performance of support vector machine classification. The same 844 marker proteins were used for all three maps. F1 scores are the harmonic mean of recall (true positives / [true positives + false negatives]) and precision (true positives / (true positives + false positives]), and were determined with leave-one-out cross validation. **D)** Number of organellar assignments, by confidence class - (I) high, (II) medium, (III) low. The 844 marker proteins are not included. **E)** Stacked profile scatter within stable protein complexes. Only complexes with at least five subunits quantified across all datasets were included. Each bar slice represents the average absolute distance to the median complex profile in one map replicate. Thinner slices reflect higher quantification precision. **F)** Inter-replicate scatter, for the 2,651 proteins profiled across all conditions and replicates. X-axis shows the average absolute distance of replicates to the corresponding average protein profile. The 70^th^ percentile values for all included profiles are: DDA: 0.080; direct DIA: 0.061; library DIA: 0.052. Lower scatter reflects higher map reproducibility (X-axis cut at 0.3; <1% of profiles not shown).

### High-throughput liquid chromatography enables faster and deeper DIA-DOMs

Based on the outstanding depth of DIA-DOMs with 100 min LC gradients, we next evaluated the performance of shorter LC formats, with the aim to establish a fast organellar mapping workflow. We used the Evosep One LC system [34], which runs pre-mixed gradients with standard lengths of 21 or 44 minutes, and reduces overhead time between samples to a few minutes. To avoid any confounding effects caused by peptide library generation, we first gauged the performance with DIA processing using the *in-silico* library (discovery DIA). We compared DIA-DOMs run with 100 min (nanoLC), 44 min or 21 min gradients (Evosep), which revealed an almost linear relationship between the number of profiled proteins and runtime (Fig. 3A and Supplementary Fig. 3A, B).

**Fig. 3.**
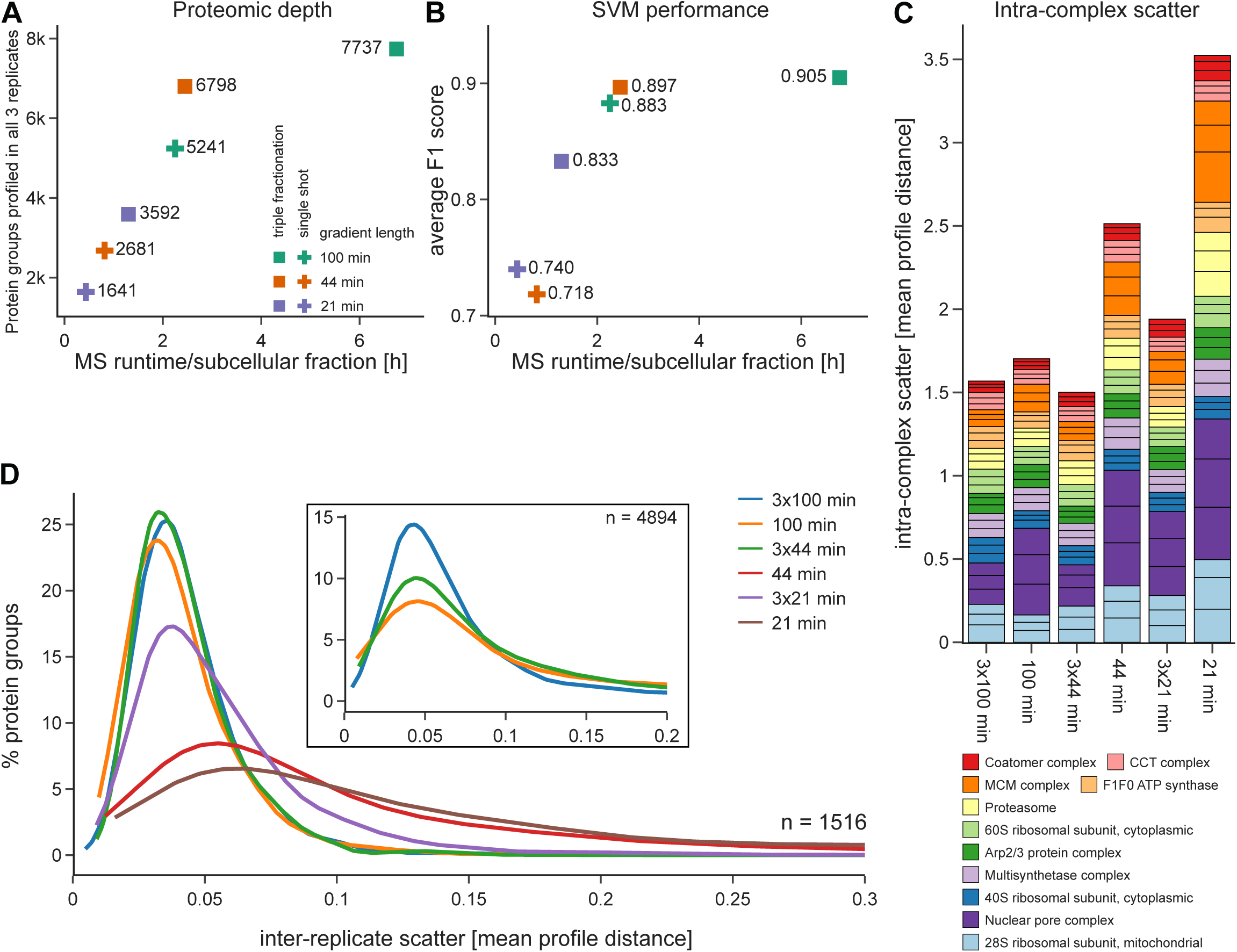
Comparison of DIA-DOMs performance with different LC gradients and sample fractionation. Comparison of maps measured with a 100 min nanoLC gradient, or with 44 min / 21 min gradients on the Evosep One LC system. SDB-RPS STAGE-tipping was employed for triple-fractionation. All maps were analyzed with discovery DIA. **A)** Proteomic depth after filtering for profile completeness across three replicates. **B)** SVM classification performance (average F1 scores). For the single-shot 100 min and the triply-fractionated 44 min and 100 min maps, the same 982 organellar marker proteins were used. For short gradient single-shot maps and the triply-fractionated 21 min maps, a smaller number of detectable marker proteins were used (21 min: 564; 44 min: 782; 3×21 min: 879). **C)** Stacked intra-complex scatter, quantified by the average absolute distance to the median complex profile. Only complexes with at least five subunits quantified across all datasets were included. Bar slices represent scatter in individual replicate measurements. **D)** Inter-replicate scatter for the 1,516 proteins quantified across all datasets. This number was limited by the depth of the 21 min dataset. (X-axis cut at 0.3; <8% of profiles not shown.) Inset: equivalent plot for the 4,894 proteins quantified across the 100 min, 3 × 44 min, and 3 × 100 min datasets, revealing the superior performance obtained with triple fractionation.

Remarkably, the 44 min gradient DIA-DOMs had a similar depth (2,681 PGs profiled across all three map replicates) as our previous 100 min gradient DDA-DOMs (Fig. 2B), and even with 21 min gradients, more than 1,600 PGs were profiled across three replicates. The organellar prediction performance of both short gradient maps was lower (Fig. 3B), but still fairly high in absolute terms (F1 = 0.7). The difference was mostly caused by substantial drops in the classifications of three organelles, Golgi, ERGIC/cis-Golgi and peroxisomes, which are particularly challenging to resolve in HeLa cells (Supplementary Fig. 3C). As expected, shortening the LC gradients also reduced profiling precision (Fig. 3C) and reproducibility (Fig. 3D). Nevertheless, even the shortest (21 min) gradient provided astonishingly well-resolved maps in only around 2.5 hours of machine time, equivalent to a throughput of over 9 maps per day. Furthermore, processing with a measured peptide library increased the depth of 21 min gradient maps to 3,105 PGs (+89%), and the depth of 44 min gradient maps to 4,411 PGs (+65%; Fig. S3D), with substantial gains in reproducibility (Supplementary Fig. 3E), making fast DIA-DOMs even more useful for high-throughput screens and rapid pilot experiments.

MaxQuant 2.0 allows DIA processing of fractionated samples, and we next explored if off-line peptide fractionation and analysis spread over several short LC runs may improve the performance relative to a ‘single-shot’ LC run of equivalent length. Since the Evosep LC system minimizes sample loading overheads, this approach also optimizes the machine-time to gradient-time ratio. We triple-fractionated our benchmark samples by peptide STAGE-tipping [25] and analyzed them with 3 × 44 min LC gradients (with 5 min overheads), which requires little more overall machine time than a single run with a 100 min nanoLC gradient (with 35 min overheads). Remarkably, fractionation yielded approximately 1,500 additional profiled PGs relative to the single shot 100 min gradient (6798 vs 5241 PGs, +30%; Fig. 3A), with similar SVM performance, precision and profile reproducibility (Fig. 3B-D).

We also analyzed maps with 3 × 21 min gradients. Relative to the 100 min gradient, this reduced machine time by approximately 50%, but incurred only a moderate drop in performance (Fig. 3A-D), and is thus a suitable intermediate format. Finally, we ran fractionated samples with 3 × 100 min nanoLC gradients, to provide ultra-deep coverage. Compared to the single-shot 100 min gradient, this yielded a further 2,496 proteins (7,737, +48%), with further improvements to SVM performance (F1 0.90 vs 0.88) and profile reproducibility (Fig. 3D). Of note, these are the deepest organellar maps from HeLa cells to date, and we provide the SVM protein subcellular localization predictions in Supplementary Table 1.

In conclusion, our data show that deep, high-accuracy DIA-DOMs can be prepared with short LC gradients, enabling high-throughput spatial proteomics. The Evosep One LC system allows convenient scaling of the runtime to the experimental requirements. In conjunction with the ability of MaxQuant to combine off-line fractionated DIA samples, single long gradients can be replaced with multiple short gradients, which enhances map performance even further.

### DIA-DOMs reveal cellular effects of starvation and Bafilomycin A1 treatment

We next tested the capabilities of DIA-DOMs for detecting induced subcellular localization changes. Nutrient deprivation in combination with Bafilomycin A1 (BafA) treatment is a widely-used method to investigate autophagy [35]. While the starvation induces metabolic changes including autophagy [36], the BafA treatment increases endo-lysosomal pH by inhibiting vATPase function and thus prevents lysosomal protein degradation [37,38]. This helps to gauge autophagic flux, and facilitates the capture of autophagic structures by imaging [35]. However, the endosome is a major protein trafficking hub, and increasing lumenal pH blocks endosomal exit pathways. As a result, proteins that normally cycle between endosomes and the plasma membrane, or between endosomes and the Golgi, become trapped in endosomes [39,40]. This ‘side effect’ of BafA treatment is largely ignored in investigations of autophagy, but may have considerable bearings on the interpretation of results. Moreover, it is not generally known which proteins get trapped, as only relatively few have been identified to date [15,39–41]. Here, we applied DIA-DOMs for the first global analysis of subcellular localization changes induced by starvation and BafA treatment.

We prepared triplicate DIA-DOMs (100 min gradients, library DIA processing) and full proteomes from HeLa cells that were either starved for 1h in the presence of BafA, or left untreated. Evaluation with DOM-QC showed that the two conditions yielded topologically very similar maps (Fig. 4A). The strength of DIA-DOMs was highlighted again by the remarkable profiling depth (>6,500 PGs in each condition), the almost complete overlap of profiled proteins (Fig. 4B) and the near identical quality (Fig. 4C) of the maps.

**Fig. 4.**
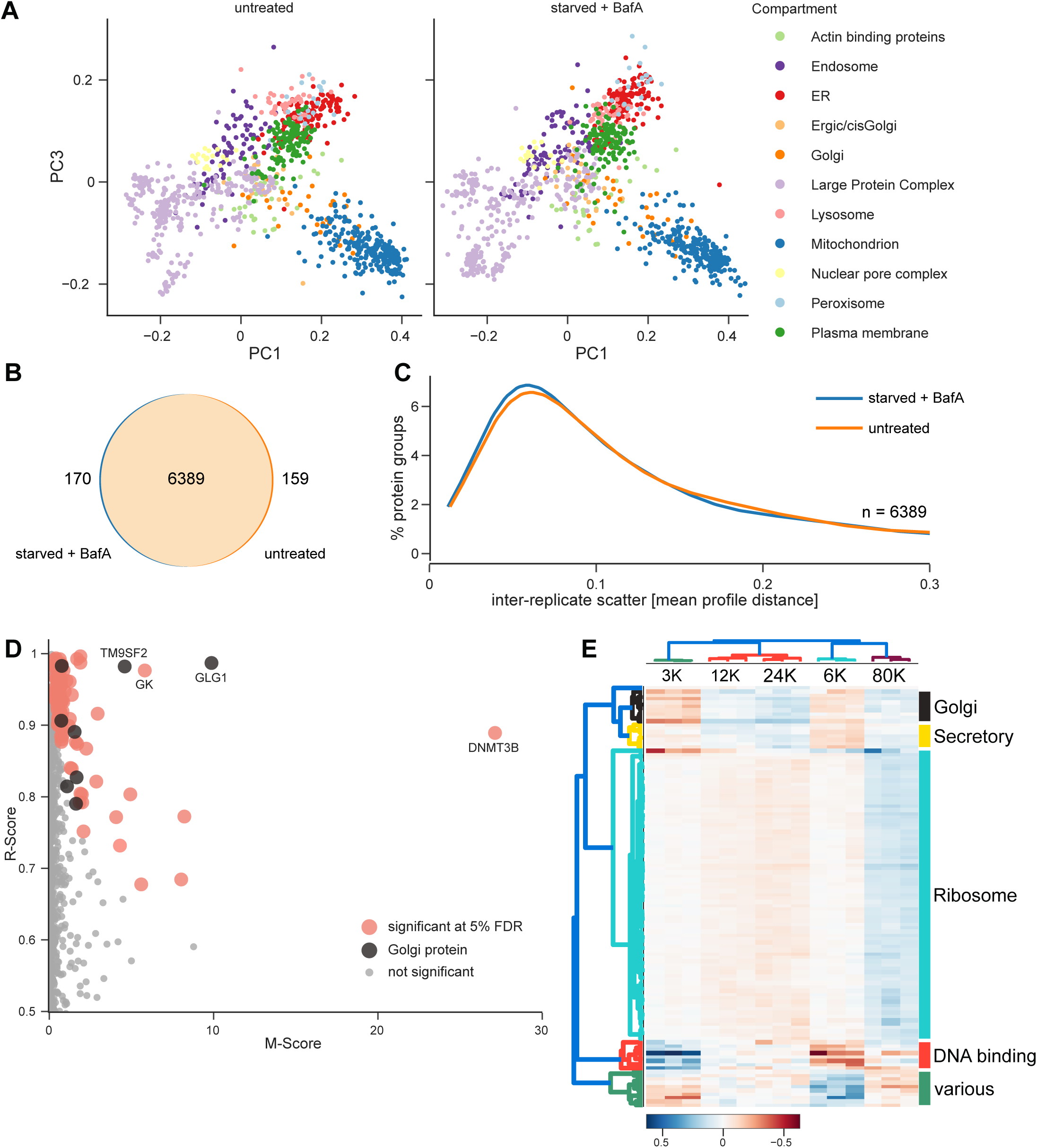
DIA-DOMs reveal the cellular effects of starvation and Bafilomycin A1 treatment. HeLa cells were either left untreated, or starved in the presence of 100 nM BafA for 1 h. DIA-DOMs were prepared in triplicate. **A)** PCA maps of organellar marker proteins show similar topology in both conditions. **B)** Overlap of profiled proteins between untreated and starved + BafA-treated maps, after filtering for profile completeness across three replicates. **C)** Inter-replicate scatter for the 6,389 proteins profiled across both conditions. **D)** Movement-Reproducibility (M-R) analysis detects 114 proteins with significantly altered subcellular localization (marked in red and black; FDR=5%). Golgi proteins with significant M-R scores are marked in black. **E)** Hierarchical clustering of the 114 significant delta profiles (starved/BafA minus control) by Pearson correlation (colour scale indicates profile change in each fraction). Cluster annotation is based on GO-term annotation enrichment (see Supplementary Fig. 4C). As expected, we did not observe the re-localization of core autophagic machinery, since HeLa cells already have high basal levels of autophagy (see Supplementary Fig. 4D).

To identify proteins with altered subcellular localization, we performed our previously established movement and reproducibility (MR) analysis [11,42]. Since starvation and BafA treatment drastically reprograms cellular metabolism, we expected pleiotropic subcellular rearrangements, in addition to endosomal trapping. Setting a high stringency cut-off (False Discovery Rate (FDR) = 5%), we identified 114 proteins with significant localization shifts (Fig. 4D; Supplementary Table 2). Of these, 103 were also quantified in our full proteomes (Supplementary Fig. 4A), and only one changed significantly in abundance. Thus, our MR analysis specifically revealed proteins that respond to starvation and BafA1 treatment by subcellular re-localization. To categorize hits functionally, we performed hierarchical clustering on the profile changes and identified five main groups (Fig. 4E; see Supplementary Fig. 4B for detailed clustering and Supplementary Fig. 4C for enrichment analysis). The first cluster contained eight Golgi proteins, while the second cluster included seven secreted/secretory pathway proteins. These proteins all shifted towards an endosomal profile, consistent with endosomal trapping (Supplementary Table 2). The third and largest group predominantly contained proteins involved in translation and mRNA processing (including over 40 core ribosomal proteins), which shifted towards the highest speed fraction (Fig. 4E and Supplementary Table 2). This is consistent with a starvation-induced reduction of translational activity, resulting in an increased number of free ribosomes or smaller translational assemblies. The remaining two clusters were more heterogeneous, and included several DNA binding proteins/transcriptional regulators. Of note, they contained two of our top hits, DNMT3B and Glycerol kinase (GK). DNMT3B is a DNA methyl transferase. Interestingly, its close homologue DNMT3A has recently been shown to exert a ‘memory’ function in autophagy, by long-term regulation of expression of autophagy-associated genes upon starvation [43], suggesting a related function for DNMT3B. GK is a key modulator of energy and lipid metabolism; its association with the outer mitochondrial membrane is regulated by starvation [44], and this response is captured by our maps (Supplementary Table 2).

The hits in Fig. 4D/E represent only the most prominent translocation events in our dataset. Therefore, we created an interactive database for the exploration of individual profile shifts (Supplementary Table 2).

Taken together, our DIA-DOMs analysis revealed a broad spectrum of known and novel subcellular rearrangements related to gene regulation and metabolism induced by starvation and BafA1 treatment, as well as endosomal trapping of diverse endomembrane proteins. Intriguingly, we identified eight Golgi proteins that shifted towards endosomes. This group contains some of the strongest hits (e.g., GLG1 and TM9SF2, Fig.4D), prompting us to characterize the behaviour of Golgi proteins in more detail.

### Systematic identification of Bafilomycin A1 sensitive anterograde cycling Golgi proteins

Endosomal trapping of some Golgi proteins has previously been reported [15,39–41], but it has not been studied systematically which Golgi proteins are susceptible. Such an analysis could distinguish Golgi proteins with anterograde cycling from those that are relatively static, which would shed light on a fundamental feature of Golgi homeostasis (Fig. 5A). Intriguingly, the eight endosome-trapped Golgi proteins identified above (Fig. 4E and Supplementary Fig. 4B) differed considerably in shift magnitudes (Fig. 4D), indicating differential degrees of trapping. This also suggested that there may be further Golgi proteins with partial shifts below the detection limit of the MR analysis at 5% FDR. For a comprehensive and systematic characterization of Golgi protein behaviour, we therefore developed a targeted cross-correlation analysis strategy. We first compiled a list of all transmembrane (32) and lumenal (2) Golgi proteins from our untreated maps. We then calculated all pairwise correlations of their shift profiles, and performed hierarchical clustering. Strikingly, two clearly segregated groups emerged from the data (Fig. 5B). The first group (Cluster 1) contained all eight proteins identified by MR analysis, which formed a particularly well-defined core. The first group also included the two lumenal Golgi proteins (SDF4 and FAM3C), suggesting that these may be subject to endosomal trapping too. To characterize cluster behaviour in detail, we plotted the shifts for all proteins in PCA space (Fig. 5C, D). Cluster 1 proteins showed largely parallel shifts of different magnitudes from Golgi to endosomes/lysosomes (Fig. 5C). In contrast, Cluster 2 proteins showed variable shifts within the Golgi boundaries (Fig. 5D). Thus, our data reveal two classes of Golgi proteins – those that undergo partial or complete endosomal trapping (Cluster 1), and those that do not show any trapping under the experimental conditions (Cluster 2). To assess the individual degree of trapping for proteins in Cluster 1, we ranked them by absolute profile shift magnitudes (Fig. 5E). The top eight proteins corresponded to the hits from our MR analysis (primary hits). Importantly, the 13 secondary hits included both GOLM1 and GOLIM4, which have previously been shown to undergo endosomal trapping [41]. Inspection of individual profile shifts allowed us to further classify the transitions as complete, partial or marginal translocations (Supplementary Table 2; we also removed three proteins with negligible shifts (YIPF3, ACSL4, IMPAD1)). Shift differences may reflect different cycling kinetics, with fast cycling proteins showing more complete transitions. For further orthogonal validation of our predictions, we selected three proteins with different profiling behaviour for imaging: GLG1 and TGOLN2 (also known as TGN46) from Cluster 1, and GALNT2 from Cluster 2 (Fig. 5F). HeLa cells were starved and BafA treated as before, or left untreated. Colocalization analysis by widefield microscopy fully confirmed our profile analysis: as predicted, GLG1 showed complete transition from Golgi to a punctate endosomal pattern, TGOLN2 underwent a partial translocation, and GALNT2 retained its Golgi pattern (Fig. 5G). Furthermore, we confirmed that the re-localization was caused by the BafA treatment, and not by starvation (Supplementary Fig. 5A, B).

**Fig. 5.**
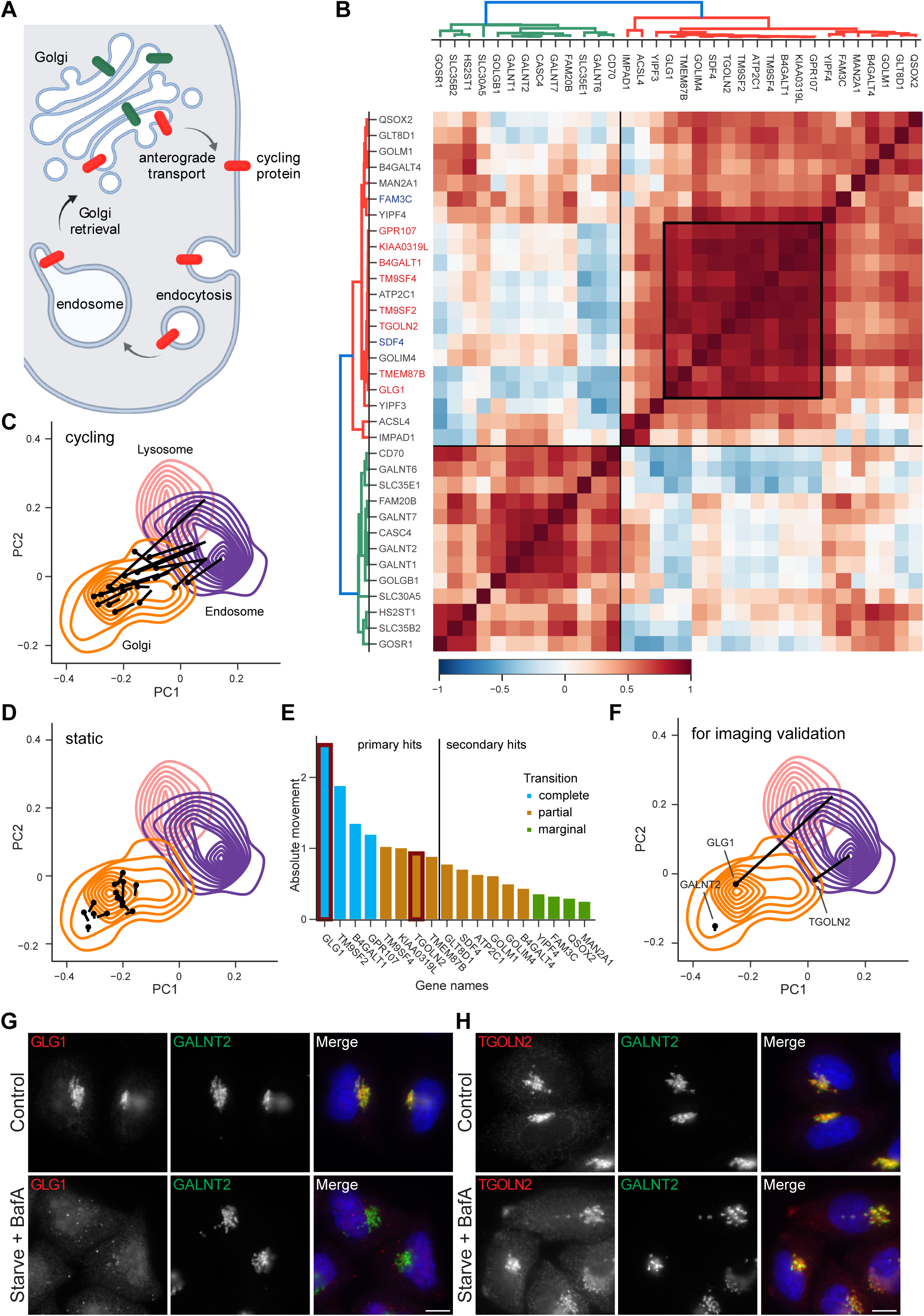
In-depth analysis and validation of Golgi protein behaviour caused by starvation and Bafilomycin A1 treatment. **A)** Some Golgi proteins cycle to the plasma membrane and back, via endosomes (red). BafA treatment compromises the retrieval pathway, trapping Golgi proteins in endosomes; non-cycling Golgi proteins (green) remain unaffected. **B)** Cross-correlation matrix and clustering of integral membrane and lumenal Golgi proteins (colour scale indicates Pearson correlation of delta profiles). Gene name colors: red = primary hits from outlier analysis, blue = lumenal proteins. Box highlights the core cluster of cycling proteins. **C)** Absolute movement distance summed over three replicates stratifies phenotype strength. The plot also reflects the sensitivity boundary of the MR outlier test (primary hits), and shows additional hits identified through the targeted correlation analysis (secondary hits). Proteins validated by imaging are highlighted with red boxes. Categorization into complete, partial and marginal shifts was based on manual evaluation of profiles. **D-F)** Protein shift trajectories starting at untreated positions overlaid with density plots of relevant organellar marker proteins. **D)** Anterograde cycling Golgi proteins (Cluster 1) shift towards the endosome/lysosome. **E)** Static Golgi proteins (Cluster 2) stay within the core density of the Golgi apparatus. **F)** Movements of three proteins selected for imaging validation. **G-H)** Widefield imaging of immunofluorescence labelling of GLG1, TGOLN2 (TGN46), and GALNT2 validated the localization shift predictions shown in **F**. HeLa cells were left untreated (control) or were starved in the presence of 100 nM BafA for 1 h. In the merged images, DAPI labelling of the nucleus is also shown (blue). Scale bars: 10 μm. Images are representative of at least 13 images per condition. **G)** GLG1 (red) completely disperses from the Golgi; **H)** TGOLN2 (red) shifts away from the Golgi; **G-H)** GALNT2 (green) remains unchanged. Note, the relocalization effects were confirmed to be caused by BafA treatment, and not by starvation, by imaging of cells that were starved without BafA or treated with BafA in full medium (see Supplementary Fig. 5A and B).

In sum, our targeted correlation-based analysis increases the sensitivity of Dynamic Organellar Maps for detecting small, but highly correlated shifts, and enables prediction of relative phenotype strength. Our systematic assessment of Golgi protein anterograde cycling behaviour illuminates an important aspect of Golgi organelle homeostasis, and demonstrates the power of DIA-DOMs for functional investigations.

## Discussion

Dynamic organellar maps (DOMs) capture protein localizations and their changes at the proteomic scale, and the approach has driven diverse discoveries in cell and medical biology [11,13–17]. The bottleneck of our original DDA-based workflow was the considerable MS time required to achieve deep coverage. Here, we introduce label-free DIA-DOMs, which overcome this limitation. DIA-DOMs achieve twice the proteomic depth of DDA-DOMs in the same MS time, and require only one third of the MS time to reach the same depth.

Three tools were instrumental for establishing DIA-DOMs. First, we created DOM-QC (https://domqc.bornerlab.org), an open source web app for the analysis of profiling data. While previous quality assessment tools focused on organellar resolution (MetaMass and QSep [45,46]), DOM-QC additionally assesses profile precision and reproducibility, which are key parameters for comparative experiments. DOM-QC enables comprehensive, standardized formatting and benchmarking of profiling data, and facilitates objective method optimization. Second, the high-throughput Evosep One HPLC system allowed us to explore short-gradient maps. While DIA-DOMs work well with a conventional nano LC, the Evosep greatly enhances MS runtime efficiency and scalability of the method. Third, MaxQuant 2.0 can process DIA samples split over several fractions, a powerful feature currently not offered by other mainstream DIA software. This allowed us to combine several long LC runs into an ultra-deep mapping analysis. Moreover, in conjunction with the Evosep, it enabled us to replace single long LC gradients with multiple short Evosep gradients of equivalent overall run time, which considerably improved performance. Based on our extensive optimization, we now recommend the following formats for DIA organellar mapping: 1) Deep mapping with 12 hours of MS time per map, either using single runs on a 100 min nanoLC gradient, or fractionation/triple runs with 44 min Evosep gradients; 2) High-throughput maps with approximately 2.5 hours MS time per map, using Evosep 21 min gradients. Suitable peptide libraries generated by additional DDA measurements further boost the performance of either format.

DIA-DOMs now replace our previous label-free DDA-DOMs [19], as they strongly improve map performance across all metrics. Our original DDA-SILAC DOMs [11,19] still have better precision than label-free DIA-DOMs (Supplementary Fig. 5C), but DIA-DOMs have similar reproducibility and offer three times greater proteomic depth in the equivalent MS run time (Supplementary Fig. 5D, E). In systems that allow metabolic labelling, DDA-SILAC DOMs may still offer an advantage where there is a need to detect very small protein translocations. Future studies should investigate if DIA-SILAC DOMs may further enhance the performance.

By achieving highly reproducible organellar profiles, the original DOMs approach enabled MS-based comparative spatial proteomics for the first time [11]. Today, several global organellar profiling approaches are firmly established, including LOPIT, PCP, SUBCELLBARCODES and DOMs [3–10]. All provide high-quality organellar maps, and have individual advantages [3,4]. DOMs have the simplest workflow, require the least MS runtime, and provide robust maps from label-free samples. We would hence argue that DIA-DOMs currently offer the easiest option for labs venturing into spatial proteomics. Our extensive protocols [42], in conjunction with our DOM-QC tool, further facilitate rapid method establishment. Of note, the Olsen lab recently introduced a fast spatial proteomics method based on chemical fractionation [9], which provides complementary insights to our centrifugation-based profiling of intact organelles. Their approach also utilizes 21 min Evosep gradients, as implemented here for rapid DIA-DOMs, and achieved a similar depth to our method, suggesting that this is a robust LC-MS approach with consistent performance across sites.

To test DIA-DOMs for phenotype discovery, we assessed protein localization changes upon nutrient starvation in the presence of Bafilomycin A1. This treatment is routinely used to investigate autophagy, but also blocks protein exit from endosomes, which causes a poorly characterized traffic jam in the endomembrane system. Our analysis mapped hundreds of protein localization changes associated with starvation and metabolic reprogramming, as well as extensive endosomal trapping of secretory pathway proteins, which can be explored through our interactive database (Supplementary Table 2). Intriguingly, a large proportion of the endosomally trapped proteins normally reside in the Golgi. To further dissect how Golgi homeostasis is affected, we performed a targeted profile shift analysis of all Golgi proteins captured by our maps. This revealed two populations: those that undergo endosomal trapping and those with persistent Golgi localization. This observation is consistent with a model in which some Golgi proteins undergo anterograde cycling via endosomes, and others do not, as previously proposed [39]. Our data now substantially expand this model. First, we provide a systematically-derived compendium of anterograde cycling Golgi proteins. Furthermore, we observe pronounced differences in the degree of endosomal trapping within the experimental timeframe. Phenotypic strength varied from complete transitions (e.g., GLG1), to partial (TGOLN2/TGN46) and very subtle shifts (e.g., GOLIM4). The simplest explanation is that proteins cycle with different kinetics and that fast-cycling proteins undergo more complete shifts. While our data support the existence of a distinct pool of static proteins, these may also cycle, but with very slow kinetics or under different physiological conditions; alternatively, the cycling route may bypass the endosome. While we currently cannot distinguish between these scenarios, our identification of large sets of apparently static and cycling proteins, and the prediction of their relative cycling speeds, will facilitate future investigations into this fundamental property of Golgi proteins.

In conclusion, DIA-DOMs enable label-free organellar profiling with unprecedented depth, speed and precision, and provide a powerful tool for systematic phenotype discovery.

## Author contributions

GB, JS and VA devised the study; VA and JS implemented the quality control tool; JS, VA and GB analyzed the data; VA and AD performed the organellar mapping experiments; AD performed the fluorescence imaging; VA ran MS acquisitions and raw data analyses; PS contributed to raw data processing in MaxQuant; JS wrote the initial draft of the manuscript; AD and GB edited the manuscript; GB supervised the project.

## Acknowledgements

We wish to thank Matthias Mann for his continued generous support. We also want to thank members of the department for Proteomics and Signal Transduction for fruitful discussions and providing starting points for the DIA method optimization: Isabell Bludau, Sophia Steigerwald, Maximilian Zwiebel, Marvin Thielert, Patricia Skowronek, Jakob Bader, Florian Meier. We also thank Jürgen Cox for providing us with MaxQuant 2.0 prior to its release. We thank the MPIB Imaging Facility for their excellent technical support. We are very grateful to Igor Paron and the column team for outstanding technical support. Some graphical elements in Fig. 1 were previously published [11], and we would like to thank Lisa Schweizer and Sophia Steigerwald for sharing their drawings of the HPLCs and the mass spectrometer.

## Materials and Methods

### Experimental Protocols Antibodies

The following antibodies were used in this study: rabbit anti-GLG1 1:200 for IF (Sigma Aldrich Cat# SAB1303679), mouse anti-GALNT2 1:200 for IF (BioLegend Cat# 682302, RRID:AB_2566611), mouse anti-LC3B 1:400 for IF (MBL International Cat# M152-3, RRID:AB_1279144), and sheep anti-TGN46 (TGOLN2) 1:200 for IF (Bio-Rad Cat# AHP500, RRID:AB_324049). Fluorescently labelled secondary antibodies were purchased from Thermo Fisher Scientific and used at 1:500 for IF: Alexa Fluor 488-labelled donkey anti-mouse IgG (Cat# A-21202, RRID: AB_141607), Alexa Fluor 568-labelled donkey anti-rabbit IgG (Cat# A10042, RRID:AB_2534017), and Alexa Fluor 680-labelled donkey anti-sheep IgG (Cat# A-21102, RRID:AB_2535755).

### Cell culture

HeLa cells [47] were cultured in Dulbecco’s Modified Eagle’s Medium (DMEM; Gibco Cat# 31966-021), supplemented with 10% (v/v) foetal bovine serum (FBS; Gibco Cat# 10270106) and 1% (v/v) penicillin-streptomycin solution (Gibco Cat# 15140122). Cells were maintained at 37 °C in a humidified atmosphere of 5% CO_2_.

### Starvation and BafA treatment

For starvation, HeLa cells were washed three times with Dulbecco’s Phosphate Buffered Saline (PBS) (Gibco Cat# 14190-094) and then incubated for 1 h in Earle’s Balanced Salt Solution (EBSS; Sigma-Aldrich Cat# E2888). Where indicated, cells were incubated in 100 nM Bafilomycin A1 (BafA, Merck, Cat# 19-148) for 1 h, in full medium (DMEM + 10% FBS) or EBSS (starve + BafA).

### Immunofluorescence microscopy

For widefield microscopy, HeLa cells were grown onto 13 mm coverslips and fixed in 3% (v/v) formaldehyde in PBS for 20 min at room temperature. Residual aldehyde groups were quenched with 20 mM glycine in PBS for 5 min. Formaldehyde fixed cells were permeabilized with 0.1% (w/v) saponin in PBS for 10 min and blocked in 1% (w/v) BSA/0.01% (w/v) saponin in PBS (BSA block) for 10 min. Primary antibody (diluted in BSA block) was added for 1 h at room temperature. Coverslips were washed three times in BSA block and then fluorophore-conjugated secondary antibody (diluted in BSA block) was added for 30 min at room temperature. Coverslips were then washed three times in PBS. Nuclei were stained with DAPI (300 nM in PBS; Thermo Scientific Cat# 62248) for 5 min. Coverslips were washed in PBS, followed by a final wash in ddH_2_O, before being mounted in ProLong™ Glass Antifade Mountant (Invitrogen Cat# P36980).

Microscopy was performed at the Imaging Facility of Max Planck Institute of Biochemistry, Martinsried, using a Leica DMi8 inverted microscope (Leica Thunder) equipped with a Leica DFC9000 GTC Camera, a 63x/1.47 oil objective (HC PL APO 63x/1.47 OIL) and an iTK LMT200 motorized stage, and controlled by Leica Application Software X (LAS X) version 3.5.5.19976. Images are representative of two biological replicates (independent starvation/BafA treatments performed on separate days), with immunofluorescence labelling and microscopy performed independently for each replicate. Cells were selected for imaging using the DAPI channel only in the Navigator software module of LAS X. ImageJ version 1.53q was used for cropping and global brightness/contrast adjustments, which were performed uniformly across all images.

### Generation of label-free organellar maps

To avoid sample-related variation during method optimization, we generated three large-scale replicate maps from HeLa cells, each from 3 × 15 cm dishes at 70-90% confluency, on a single day. Protein samples were digested with LysC and Trypsin (see below). All subsequent peptide clean-ups and peptide fractionations were performed from the same set of digests. For the comparative experiment (Fig. 4 and 5), organellar maps were prepared from control HeLa cells (untreated) and HeLa cells that had been starved for 1 h in the presence of Bafilomycin A1 (100 nM), in triplicate, each from 1 × 15 cm dish at 70-90% confluency. All six maps were generated on the same day.

### Dynamic Organellar Maps workflow

Cell lysis and subcellular fractionation were performed as reported previously [11,42]. All steps were performed at 4 °C with pre-chilled ice-cold buffers. HeLa cells were washed in PBS (without CaCl_2_ and MgCl_2_), incubated in PBS for 5 min, rinsed with hypotonic buffer (25 mM Tris·HCl, pH 7.5, 50 mM sucrose, 0.5 mM MgCl_2_, 0.2 mM EGTA), and immediately incubated in hypotonic buffer for 5 min. Cells were drained, scraped into a total volume of 4 mL of fresh hypotonic lysis buffer and mechanically lysed with 15 strokes of a pre-chilled Dounce homogenizer (7 mL, tight pestle, Kontes Glass Co.). Sucrose was restored to 250 mM with hypertonic sucrose buffer (25 mM Tris·HCl, pH 7.5, 2.5 M sucrose, 0.5 mM MgCl_2_, 0.2 mM EGTA).

All centrifugation steps were performed at 4 °C with the fastest acceleration and deceleration settings. Crude cell lysates were centrifuged at 1000 × g for 10 min (Multifuge 1 L, Heraeus) to pellet nuclear material and unbroken cells (‘1K’ fraction). Post-nuclear supernatants were transferred to new tubes and centrifuged at 3,000 × g for 10 min (‘3K’ fraction). Post-3000 × g supernatants were transferred to ultracentrifuge tubes and further sub-fractionated using the Optima™ MAX Ultracentrifuge (Beckman Coulter) with a pre-chilled TLA 110 rotor (Beckman Coulter) by sequential centrifugation steps, each time collecting a protein pellet and transferring the supernatant to a new ultracentrifuge tube: 10,000 rpm (5,400 × g) for 15 min (‘6K’ fraction), 15,000 rpm (12,200 × g) for 20 min (‘12K’ fraction), 21,000 rpm (24,000 × g) for 20 min (‘24K’ fraction), and 38,000 rpm (78,400 × g) for 30 min (‘80K’ fraction). All pellets were resuspended in 1×SDS buffer (2.5 % SDS, 50 mM Tris·HCl, pH 8.1). The supernatant obtained after the final centrifugation step (cytosolic fraction) was mixed at a 4:1 ratio with 5×SDS buffer (12.5 % SDS, 50 mM Tris·HCl, pH 8.1). Samples were heated at 72 °C for 5 min and sonicated using a Bioruptor (Diagenode Inc) with fifteen 30 s on/off cycles at maximum intensity. Fully solubilized samples were stored at -80 °C. Protein concentrations were determined using the Thermo Scientific™ Pierce™ BCA (bicinchoninic acid) Protein Assay Kit (Thermo Scientific™ Cat# 23225). Following concentration determination, DTT (Sigma-Aldrich Cat# D0632-25G) was added to a final concentration of 1 mM before preparing the samples for mass spectrometry.

### Sample preparation for mass spectrometry

#### In-solution digestion of proteins

Protein was precipitated by the addition of five volumes of ice-cold acetone, incubated at -20°C overnight and pelleted by centrifugation at 10,000 × g (Centrifuge 5418R, Eppendorf) for 5 min at 4°C. All subsequent steps were performed at room temperature. Precipitated protein pellets were drained, air-dried for 5 min, resuspended thoroughly in urea buffer (8 M urea, 50 mM Tris·HCl, pH 8.1, freshly added 1 mM DTT), and incubated for 15 min. Sulfhydryl groups were alkylated by the addition of 5 mM iodoacetamide for 1 h in the dark. Proteins were enzymatically predigested by the addition of LysC (1 µg per 50 µg of protein; Wako Cat# 129-02541) for overnight incubation. Predigests were then diluted four-fold with 50 mM Tris, pH 8.1 (final urea concentration < 2 M) before addition of trypsin (1 µg per 50 µg of protein; Sigma-Aldrich Cat# T6567) for a 3 h incubation. The reaction was stopped by the addition of 1 % trifluoroacetic acid (TFA, final pH < 3). Samples were incubated on ice for 10 min and spun at 10,000 × g for 5 min at 4 °C. Supernatants were transferred to new tubes for peptide storage at -20 °C.

#### Purification and fractionation of peptides

Peptides were purified either by solid-phase extraction with poly(styrenedivinylbenzene) reverse-phase sulfonate (SDB-RPS), as previously described [25], or by LC trapping using commercially available C18 StageTips (EvoTips Cat# EV2001) of the Evosep System, according to the manufacturer’s instructions. In brief, EvoTips were activated by wetting the C18 material in 1-propanol, washed with Evosep buffer B (0.1 % [v/v] formic acid in acetonitrile), and wetted in 1-propanol again for 5 min. Soaked tips were washed with Evosep buffer A (0.1 % [v/v] formic acid), then with 0.2 % formic acid and then loaded with 200 ng acidified peptide sample. EvoTips were washed with Evosep buffer A, and finally loaded with Evosep buffer A and stored at 4 °C until analysis by mass spectrometry. Peptides purified via the SDB-RPS approach were dried at 45 °C in a centrifugal vacuum concentrator (Concentrator 5301, Eppendorf), resuspended in buffer A* (0.1 % [v/v] TFA, 2 % [v/v] acetonitrile), and stored at -20 °C until analysis by mass spectrometry.

For deep measurements and the DDA library for the 100 min gradient, peptides were triple-fractionated on SDB-RPS StageTips [25]. StageTips were washed with 100% acetonitrile, equilibrated with StageTip equilibration buffer (30% [v/v] methanol, 1% [v/v] TFA), and washed with 0.2 % (v/v) TFA. 20 μg of peptides in 1 % TFA were loaded onto activated stage-tips, washed with isopropanol, and then twice with 0.2% (v/v) TFA. Peptides were eluted in three consecutive fractions by applying a step gradient of increasing acetonitrile concentrations: 20 μL SDB-RPS-1 (100 mM ammonium formate, 40 % [v/v] acetonitrile, 0.5 % [v/v] formic acid), then 20 μL SDB-RPS-2 (150 mM ammonium formate, 60 % [v/v] acetonitrile, 0.5 % [v/v] formic acid), then 30 μL SDB-RPS-3 (5 % [v/v] NH_4_OH, 80 % [v/v] acetonitrile). For the DDA libraries for the 21 and 44 min gradients, peptides were fractionated into a final eight fractions using a Pierce High pH reversed-Phase Peptide Fractionation Kit (Thermo Fisher Scientific, 84868), according to the manufacturer’s instructions.

#### Mass spectrometric analysis

All measurements were performed on a Thermo Exploris 480 mass spectrometer, with minimal chromatography column changes. Several MS setups and strategies were tested, most importantly data independent vs data dependent acquisition. The effect of gradient length on map quality was evaluated for 21, 44 and 100 min gradients, for both triply SDB-RPS fractionated [25] and unfractionated samples.

#### Liquid Chromatography

Nanoflow reversed-phase chromatography was performed using either the Evosep One (Evosep Biosystems) or the EASY-nLC 1200 ultra-high-pressure system coupled online to an Orbitrap Exploris 480 instrument via a nano-electrospray ion source (all Thermo Fisher Scientific). On the EASY-nLC 1200 system a binary buffer system with the mobile phases A (0.1 % [v/v] formic acid) and B (80 % acetonitrile, 0.1 % [v/v] formic acid) was employed. Peptides were separated in 100 min at a constant flow rate of 300 nL/min on a 50 cm × 75 µm (i.d.) column with a laser-pulled emitter tip, packed in-house with ReproSil-Pur C_18_-AQ 1.9 µm silica beads (Dr. Maisch GmbH). The column was operated at 60 °C using an in-house manufactured oven. In total, 300 ng of purified peptides in Buffer A* were loaded onto the column in Buffer A and eluted using a linear 84 min gradient of Buffer B from 5 % to 30 %, followed by an increase to 60 % B in 8 min, a further increase to 95 % B in 4 min, a constant phase at 95 % B for 4 min, followed by washout – a decrease to 5 % B in 5 min and a constant phase at 5 % B for 5 min – before re-equilibration. On the Evosep One LC system a binary buffer system with the mobile phases A (0.1 % [v/v] formic acid) and B (0.1 % [v/v] formic acid in acetonitrile) was used. Peptides were separated in 21 min at a flow rate of 1.0 µL/min on an 8 cm column (with a throughput of 60 samples per day [SPD]) or 44 min at a flow rate of 0.5 µL/min on a 15 cm column (with a throughput of 30 SPD), using in-house packed columns and standard pre-programmed gradients. The 15 cm in-house packed column was operated at 60 °C using an in-house manufactured oven.

#### Mass spectrometry

For DDA, the Orbitrap Exploris 480 mass spectrometer run by Xcalibur (v.4.4, Thermo Fisher) was operated in top 15 scan mode (DDA) with a full scan range of 300 - 1650 Th when coupled to the EASY-nLC 1200 system (100 min gradient). Survey scans were acquired at 60,000 resolution with an automatic gain control (AGC) target of 3 × 10^6^ charges and a maximum ion injection time of 25 ms. The selected precursor ions were isolated in a window of 1.4 Th, fragmented by higher-energy collisional dissociation (HCD) with normalized collision energies of 30. Fragment scans were performed at 15,000 resolution, with a maximum injection time of 28 ms, an AGC target of 1 × 10^5^ charges, and a precursor dynamic exclusion for 30 s. Acquisition schemes for the data-independent acquisition (DIA) scan mode used here were described previously [48,49], but were optimized and tailored for the Dynamic Organellar Maps approach. In brief, the DIA method for the 100 min gradient consisted of one survey scan that was followed by 33 variably sized MS2 windows (17-161 Th) in one cycle, resulting in a cycle time of 2.5 s. Survey scans were acquired at 120,000 resolution with an AGC target of 3 × 10^6^ charges and a maximum injection time of 60 ms covering a m/z range of 350 – 1,400. MS2 scans were acquired at 30,000 resolution with an Xcalibur-automated maximum injection time, covering a m/z range of 332 (lower boundary of the first window) to 1,570 (upper boundary of the 33rd window). The DIA method for the 44 min and 21 min gradient consisted of one survey scan that was followed by 35 equally sized MS2 windows (19.2 Th with 1 Th overlap) in one cycle, resulting in a cycle time of 1.5 s. Survey scans were acquired at 120,000 resolution with an AGC target of 3 × 10^6^ charges and a maximum injection time of 45 ms, covering a m/z range of 350 – 1,400. MS2 scans were acquired at 15,000 resolution with a maximum injection time of 22 ms, covering a m/z range of 361 - 1,033.

### Bioinformatic analysis

#### Raw data analysis

For peptide and protein identification, MS raw data were imported into MaxQuant version 1.6.7 or 2.0.0.0 [32] (see below). Unless otherwise stated, default parameters were used for all settings. The MS2 spectra were searched against the SwissProt entries contained in the UniProt human reference proteome FASTA database (UP000005640_9606, 42,418 entries). Spectral libraries were constructed using DDA raw data of fractionated subcellular samples of the same organellar maps that were used for the data acquired in DIA mode.

#### Spectral library generation and DDA analysis

For spectral libraries and the DDA benchmark, DDA raw files were processed in MaxQuant (v.1.6.7) [30,50] employing the Andromeda search engine [51]. For accurate label-free quantification, the ‘MaxLFQ algorithm’ [20] was enabled with LFQ minimum ratio count of 1 and the match-between-runs feature was enabled to match between equivalent and adjacent subcellular fractions of replicates. Each spectral library was assembled from 21 samples or 56 samples (six subcellular fractions as used for mapping, plus cytosol, each fractionated at the peptide level 3-fold or 8-fold as described above). A dedicated library was generated for each LC gradient length (100 min, 159,000 peptides; 44 min, 89,000 peptides; 21 min, 59,500 peptides).

#### DIA analysis

DIA raw files were processed via MaxDIA [32], which is embedded into the MaxQuant software environment (v.2.0.0.0), using default settings except for using a minimum LFQ ratio count of 1. For both the discovery and library DIA approaches, spectral libraries of peptides were provided in the form of ‘peptides’, ‘evidence’, and ‘msms’ files. Whereas for the library approach these files were obtained from MaxQuant DDA searches, for the discovery approach an *in silico* predicted library for all human peptides with up to 1 missed cleavage was used. The prediction had previously been generated using the DeepMass:Prism tool [33]. The provided library was filtered to contain only Swiss-Prot entries, using a python script (github.com/cox-labs/DIAtools/tree/main/Misc/FilterAdditional).

#### Downstream data quality analysis using the DOM-QC web app

The intra- and inter-experimental quality of the dynamic organellar maps was evaluated to assess the performance of different combinations of MS methods, LC-MS setups, and processing strategies. To enable the visual exploration and quality assessment of spatial proteomics data, we developed DOM-QC, a web-based app (https://domqc.bornerlab.org). The workflow is entirely based on the Python scripting language and uses several external libraries as documented on github (https://github.com/valbrecht/SpatialProteomicsQC). DOM-QC performs customizable data filtering, normalization, and graphical representation. Various analysis tools allow detailed exploration of data, map quality, reproducibility, and resolution. All results can be downloaded as support vector graphics, formatted tables and as comprehensive .json files for custom analysis. Importantly, several maps can be compared in parallel. With the exception of the protein movement analysis (see below), all map analyses shown in the paper were performed using six-point profiles, i.e., protein abundance across 1K, 3K, 6K, 12K, 24K and 80K fractions.

#### Data filtering

The primary output from MaxQuant or Spectronaut, or preprocessed data with protein quantification across the subcellular fractions, can be loaded into DOM-QC. For the MaxQuant output, reverse hits, contaminants and proteins only identified by modifications are removed. Further filtering is then performed at the level of individual maps and tailored to each quantification strategy, to obtain datasets with high-quality measurements.

For SILAC maps, SILAC ratios are retained if they are based on more than two quantification events, or on two quantification events where the ratio variability was below 30 %. For each fraction, SILAC ratios are normalized by dividing by the median ratio for the fraction. Only proteins with complete profiles are retained, i.e., a valid SILAC ratio in each subcellular fraction. SILAC ratios are inverted (assuming that the reference fraction is SILAC heavy [11]) and profiles for each protein are 0-1 normalized. For LFQ maps, intensities are already globally normalized, hence no further normalization is required. Two stringency filters are applied: First, only profiles with LFQ intensities in at least 4 consecutive fractions are considered. Second, profiles are rejected if their average MS/MS count per subcellular fraction is less than two. Then, (0-1) normalization of each profile is performed. Filtered datasets were annotated based on a predefined set of 1076 organellar marker proteins covering 12 subcellular localizations/organelles [11]. These default settings were used for all datasets in this study.

#### Protein group alignment

To compare quantifications from different raw data processing runs, protein groups need to be aligned. We implemented a strategy in which we prioritize matching of single-gene locus protein groups with complete coverage across experiments, over matching of (rare) multi-gene locus groups and groups with incomplete coverage. First, single-gene locus protein groups are temporarily reduced to the canonical id (if present), and otherwise to the first listed isoform id. Protein groups that can be found in all compared runs are then re-labelled to the reduced protein group id and flagged as ‘primary id’ matches. Typically, this is the case for the majority of proteins. Second, we maximize overlap for the remaining multi-gene locus groups, and single-gene locus groups with incomplete coverage, at the cost of making less exact matches. Starting with the largest multi-gene locus group across all experiments, we match this with its largest remaining subset in each other experiment, removing them from the pool of available groups. This is repeated until no multi-gene locus groups remain. These matches are re-labelled with all protein ids contained within the group, and either flagged as ‘multiple genes’, if all matched protein groups cover identical loci across experiments, or as ‘gene level conflict’, if different loci were covered in different experiments. At the end of this procedure only single-gene locus groups with incomplete coverage remain to be aligned. These are reduced in the same way as the single-gene locus full coverage groups and are also flagged as ‘primary id’ matches. To ensure full traceability of the original protein grouping in each of the compared search engine runs, an id mapping table is stored and available for download.

#### Assessment of proteomic depth

Proteomic depth is assessed by counting protein groups that are either identified in one or all replicates, and by counting proteins that are fully profiled (i.e., passing all quality filters) in one or all replicates. Venn diagrams and upsetplots are provided to evaluate overlap of proteins.

#### Principal component analysis

For graphical map representation, filtered and (0-1) normalized data from all experiments compared were centered and scaled to unit variance in each fraction, and jointly subjected to principal component analysis (PCA) to achieve dimensionality reduction. For each map, the first three principal components were calculated via Python’s scikit-learn library (v.0.23.2). Scores plots of PCs 1 and 3 usually provide the best visual resolution of clusters.

#### Profile scatter within stable complexes (intramap scatter)

The subunits of a stable protein complex have identical subcellular distributions, and should therefore have very similar abundance profiles. Observed deviations are mostly caused by MS measurement noise, and intra-complex protein scatter thus reflects within-map quantification precision. We curated a dataset of around 30 well-characterized protein complexes with at least five subunits (e.g., 20S core proteasome, CCT, COPI). Within DOM-QC, starting with the filtered and (0-1) normalized data, profiles that belong to a specified protein cluster are extracted and filtered to leave only proteins that were measured with full coverage across all compared maps and experiments. By default, only complexes with full coverage data for at least five proteins are analyzed. Subsequently, the absolute distance of each subunit profile to the complex median profile is calculated. Smaller distances suggest more precise quantification. We observed that the baseline “tightness” varies somewhat between complexes, and hence recommend that the overall assessment should be based on at least ten different complexes.

#### Inter-maps profile reproducibility (intermap scatter)

To evaluate the reproducibility of (0-1) normalized profiles, the inter-profile scatter across replicates is calculated. For each protein, the absolute distance of each replicate profile to the average profile from all replicates is calculated; these distances are averaged to obtain this protein’s profile scatter. Global profile scatter is then plotted as a density function for proteins common to all compared maps. As an additional output, the distribution for all profiled proteins can be displayed, regardless of overlap with the other examined maps. The greater the proportion of proteins with low scatter, the better the between-maps reproducibility. As a numeric readout, the scatter at a specified quantile of each distribution can be displayed.

#### Support vector machine analysis

To further evaluate the performance of organellar maps, their power to predict protein localization was assessed using quality-filtered, (0-1) normalized data with full replicate coverage. For supervised classification a set of marker proteins covering 11 subcellular localizations was used as a means to assign all other proteins to organellar clusters by SVMs in Perseus (v.1.6.2.3) [52]; the previously defined ER_high_curvature cluster [11] was removed in this study due to the low number of marker proteins in the depth-limited short-gradient datasets. As far as practicable (see figure legends), only markers present in all compared datasets were included, and identical SVM parameters were used. First, the SVM algorithm was trained on the marker proteins, to determine optimal classification parameters and thus define optimal boundaries between the organellar clusters. The kernel was set to a radial basis function (RBF) and 8-fold cross-validation was used to ensure models are not overfit. Both parameters, Sigma and C were optimized via one-dimensional scans to achieve the minimal overall classification error. Second, optimized parameters were used to classify non-marker proteins applying leave-one-out cross-validation. The misclassification matrix from Perseus was then imported into the DOM-QC tool to calculate the global marker prediction recall (proportion of correctly predicted to the total number of markers), the organelle specific recall (proportion of markers correctly assigned to the cluster), and the organelle specific precision (ratio of markers correctly assigned to the number of all markers assigned to the cluster). The harmonic mean of recall and precision, the F1 score, was used as the primary readout for SVM performance.

#### Protein movement analysis

The previously established MR plot analysis was used to identify protein profiles that are significantly shifted between treatments [11,42], with minor modifications. Here, we compared organellar maps from untreated HeLa cells and from HeLa cells that had been starved in the presence of BafA for 1h. This analysis was performed in the Perseus environment (v.1.6.2.3). Replicates were numbered 1 to 3. As before [19], the 1K fractions were omitted, and profiles were 0-1 normalized to the remaining five datapoints. Next, ‘delta profiles’ were calculated within each cognate pair of untreated and treated maps, by subtracting the profiles obtained from each treated map from the profiles of its control map. Second, for each of three sets of delta profiles, a robust multidimensional outlier test was performed providing three p-values for movement for each protein. The median of the three p-values was selected for further analysis, and -log(10) transformed to obtain the Movement score (M score). Third, the Reproducibility score (R score) was calculated: the Pearson correlation of all pairs of delta profiles was calculated (Rep 1 vs 2, 1 vs 3, 2 vs 3) and the minimum of these (worst correlation) was chosen as the R score. M and R scores were then combined into a single score, using the formula CombScore = M x R^7 (R scores were thus strongly weighted). As a further filter, we also calculated profile reproducibility as the lowest Pearson correlation of all six replicate profiles (untreated 1vs2, 1vs3, 2vs3; treated 1vs2, 1vs3, 2vs3). For proteins where profile reproducibility was lower than 0.7, the CombScore was set to 0 to effectively exclude them from the analysis. For FDR calculation, the six maps were compared in all those combinations not expected to yield relevant shifts (Treated1 vs Treated2, Treated1 vs Treated3, Treated2 vs Treated3, Not Treated1 vs Not Treated2, Not Treated1 vs Not Treated3, NotTreated2 vs Not Treated3), to obtain six largely static delta profiles as decoy distributions. From this comparison, six mock p-values and six mock correlations were calculated. To get a stringent FDR, the R score was determined as the second highest correlation of these, and the M score as the -log10 median p value. The combined score was calculated as above. All proteins were then compiled into a joint list, with two entries for each protein (CombScore mock experiment and CombScore comparative experiment). Entries were then ranked by CombScore. FDR was calculated as the proportion of entries from the mock experiment at a given CombScore cut-off. FDR thus converged onto 50% for low CombScores (equal proportion of real and mock hits). We chose an FDR of 5% for generating the list of 114 hits shown in Fig. 4.

#### DOM-QC 1-min Quick Start Guide

This guide will allow you to test the DOM-QC tool with data generated in this study.

1. Go to webpage https://domqc.bornerlab.org.
2. Click on the big green button ‘Benchmark multiple experiments’.
3. From the ‘Add reference set’ drop-down menu (top right corner), select ‘HeLa 1×100min libraryDIA’. Click the ‘Load’ button.
4. Repeat Step 3., to load the ‘HeLa 1×100min DDA’ file. You have now loaded two different sets of maps.
5. Click the big green button ‘Align and analyse selected datasets’. This may take a moment – the program will update on progress and tell you when it’s finished (bottom right corner).
6. Scroll down, and select the ‘Overview’, ‘PCA maps’, ‘Depth and coverage’,… tabs to view the different analyses.

The sample ‘reference sets’ .json files are already integrated into the DOM-QC tool. To analyse the complete datasets generated in this study, upload the .json files provided as Supplementary Data files 1-5. To do so, in step 3., click the ‘Browse’ button, select a .json file, and upload.

To configure your own analysis of profiling data, go to the start page and follow the instructions.

## Data availability

The mass spectrometry proteomics data generated in this study will be deposited to the ProteomeXchange Consortium [http://proteomecentral.proteomexchange.org] via the PRIDE partner repository, prior to publication, and will be made available to reviewers upon request.

All DOM-QC analyses in this study can be replicated by upload of the .json files, provided as Supplementary Data 1-5, to https://domqc.bornerlab.org. See DOM-QC 1-min Quick Start Guide above.

## Code availability

The web app DOM-QC introduced in this study is available at https://domqc.bornerlab.org, and the source code at https://github.com/valbrecht/SpatialProteomicsQC.

---

**Supplementary Fig. 1.**
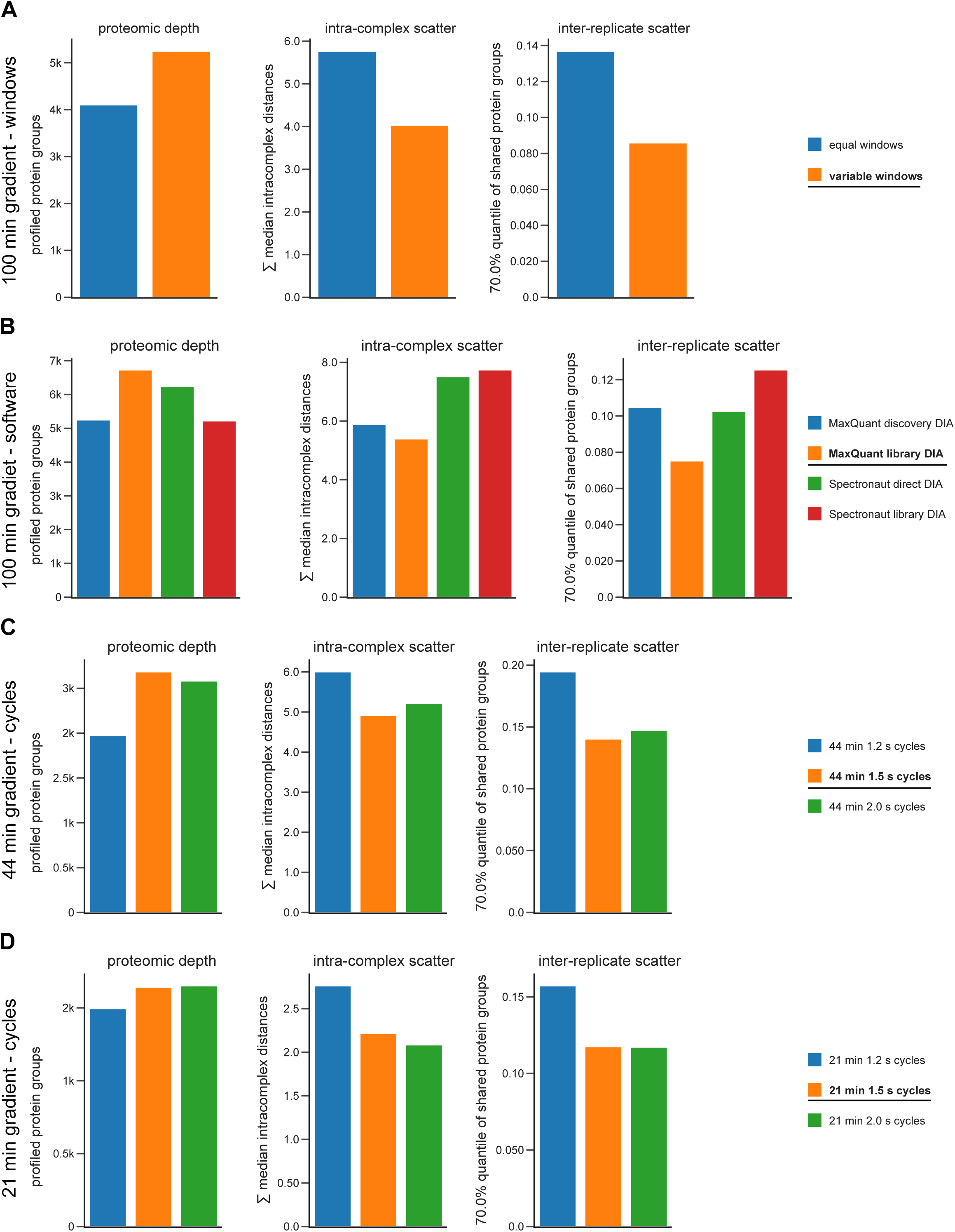
Optimization of the DIA workflow. Multiple datasets were acquired with different acquisition parameters and data processing strategies. The software app DOM-QC (https://domqc.bornerlab.org) was applied for unbiased performance evaluation. In each case, three replicate maps were analyzed, which all stemmed from the same subcellular fractionation experiment. We scored proteomic depth (number of protein groups profiled across all replicates, the higher the better), intra-complex scatter (lower scatter indicates higher quantification precision), and inter-replicate scatter (lower scatter indicates higher reproducibility). In each panel, the configuration selected for this study is highlighted in the legend (bold, underlined). **A)** Window size optimization (variable DIA windows vs equal sized DIA windows). **B)** Software (MaxQuant 2.0 vs. Spectronaut 14) and database (direct/discovery vs. library) for data processing. The same set of raw files was processed in four different ways for this comparison. MaxQuant processing yields better quantification precision and reproducibility. **C)** Cycle time optimization for the 44 min gradient. The shorter the cycle time, the more quantifications per peak (q/p) were collected; 1.2 s ∼ 7 q/p; 1.5 s ∼5.0 q/p; 2.0s ∼4 q/p. However, shorter cycle times necessitate larger DIA windows, which negatively impacts on depth and precision. **D)** Cycle time optimization for the 21 min gradient; 1.2 s ∼ 4.5 q/p; 1.5 s ∼4.0 q/p; 2.0s ∼3.5 q/p.

**Supplementary Fig. 2.**
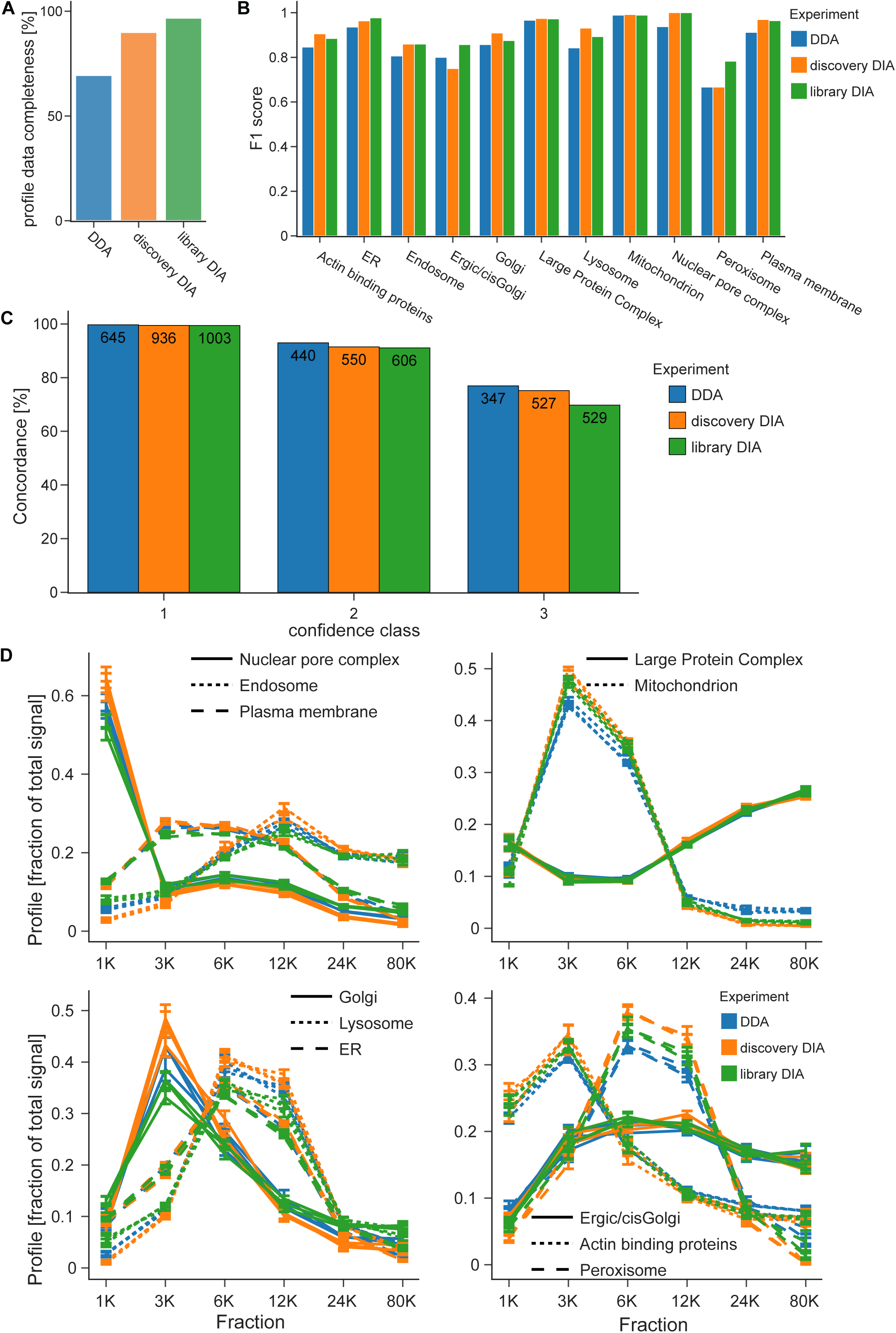
Comparison of DDA, discovery DIA and library DIA based maps, additional data. **A)** Profile data completeness of all identified protein groups, prior to quality filtering. **B)** Breakdown of SVM classification performance (F1 score) by compartment. **C)** Concordance of SVM predictions with previously published SILAC-based organellar maps [11,19], by prediction confidence class (1, high confidence; 2, medium confidence; 3, low confidence). Callouts indicate how many (non-marker) proteins in each class/experiment were shared with the SILAC reference maps; concordance indicates the proportion of agreement. **D)** Median marker profiles of classified compartments show only minor differences with the different acquisition modes.

**Supplementary Fig. 3.**
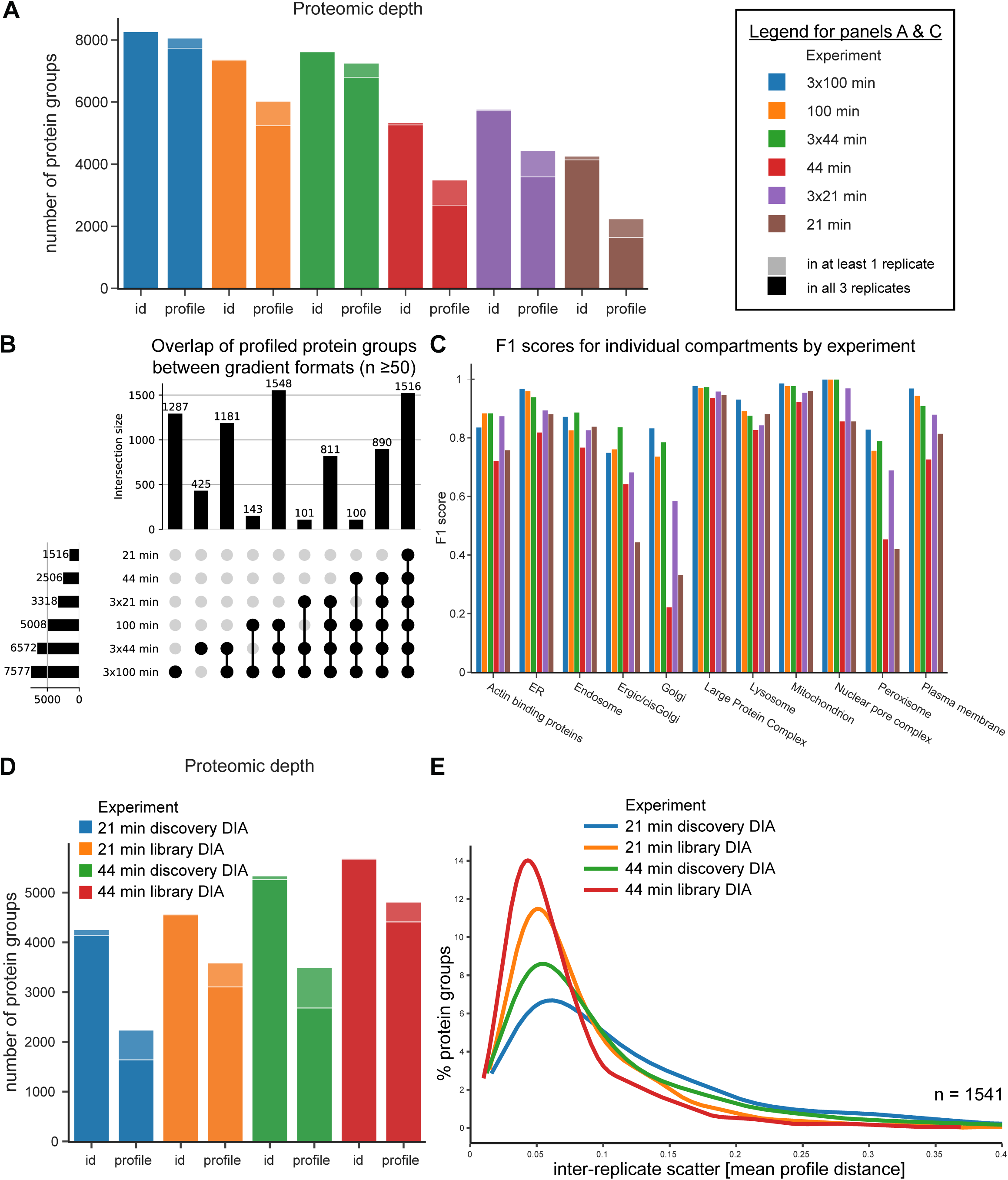
Comparison of DIA-DOMs performance with different LC gradients and sample fractionation, additional data. All datasets in **A-C** were generated with discovery DIA. **A)** Assessment of proteomic depth. id, number of protein groups identified; profile, number of protein groups with a complete abundance profile suitable for organellar mapping. Light shade, identified/profiled in at least one replicate; dark shade, identified/profiled in all three replicates. Samples were analysed either in ‘single shot’ with the indicated LC gradient time per subcellular fraction, or in ‘triple shot’ following 3 x SDB-RPS peptide fractionation. **B)** Upset-plot showing overlap of proteins profiled with different LC gradients. Combinations with intersection sizes <50 were not included. **C)** Breakdown of SVM classification performance (F1 score) by compartment and LC gradient. **D-E**, comparison of library and discovery DIA for short LC gradients. Auxiliary peptide libraries were generated by DDA, separately for each gradient (21 min gradient, 59,500 peptides; 44 min gradient, 89,000 peptides). **D)** Proteomic depth, as in **A. E)** Inter-replicate scatter, for the 1,541 proteins profiled across all experiments. Lower scatter reflects higher map reproducibility.

**Supplementary Fig. 4.**
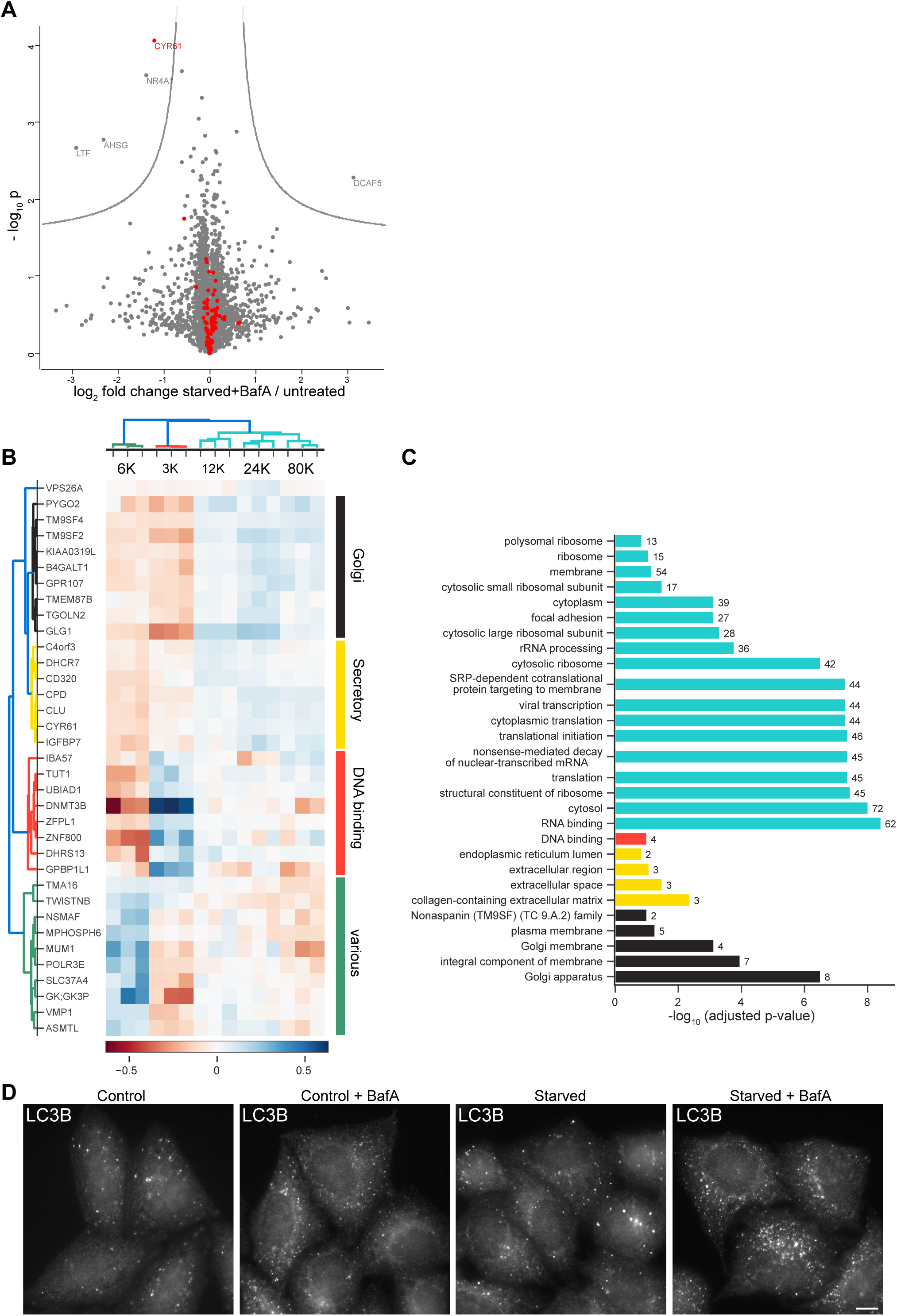
DIA-DOMs reveal the cellular effects of starvation and Bafilomycin A1 treatment, additional data. **A)** Full-proteome protein abundance analysis of HeLa cells either left untreated, or starved in the presence of BafA for 1 h. Triplicates (n=3) were analysed for each condition. 5,819 proteins were quantified in at least 5 out of 6 experiments, and included in the analysis. Data were analysed with a two-tailed *t*-test: volcano lines indicate significance thresholds (permutation-based FDR = 5%). 103 proteins with shifted subcellular localisation (as indicated by MR analysis, Fig. 4) are highlighted in red. Only one of them showed a significant abundance change. **B)** Detailed analysis of the hierarchical clustering shown in Fig. 4E. Delta profiles of proteins with a significant subcellular localisation shift were clustered by correlation, yielding five main clusters. Four clusters are shown in detail here. The fifth cluster (turquoise, not shown here) contains core ribosome proteins and translation associated proteins (see Supplementary Table 2 for a complete list). Please note that the ‘Golgi’ cluster also contains a non-Golgi protein (Pygo2), which was not included for the analysis in Fig. 5. Furthermore, VPS26A (top of the figure panel) is in a separate cluster (annotated as ‘cluster 0’ in Supplementary Table 2). **C)** Enriched GO-terms for the clusters shown in **B** and in Fig. 4E. The green cluster (annotated as ‘various’) did not show any significantly enriched terms. **D)** HeLa cells have a high level of basal autophagy. Widefield imaging of immunofluorescence labelling of LC3B as a marker for autophagosomes. Cells were either untreated (control), treated with 100 nM BafA for 1 h (control + BafA), starved for 1 h, or starved and treated with 100 nM BafA for 1 h. LC3B positive structures are readily detectable even under untreated control conditions. Scale bar: 10 µm.

**Supplementary Fig. 5.**
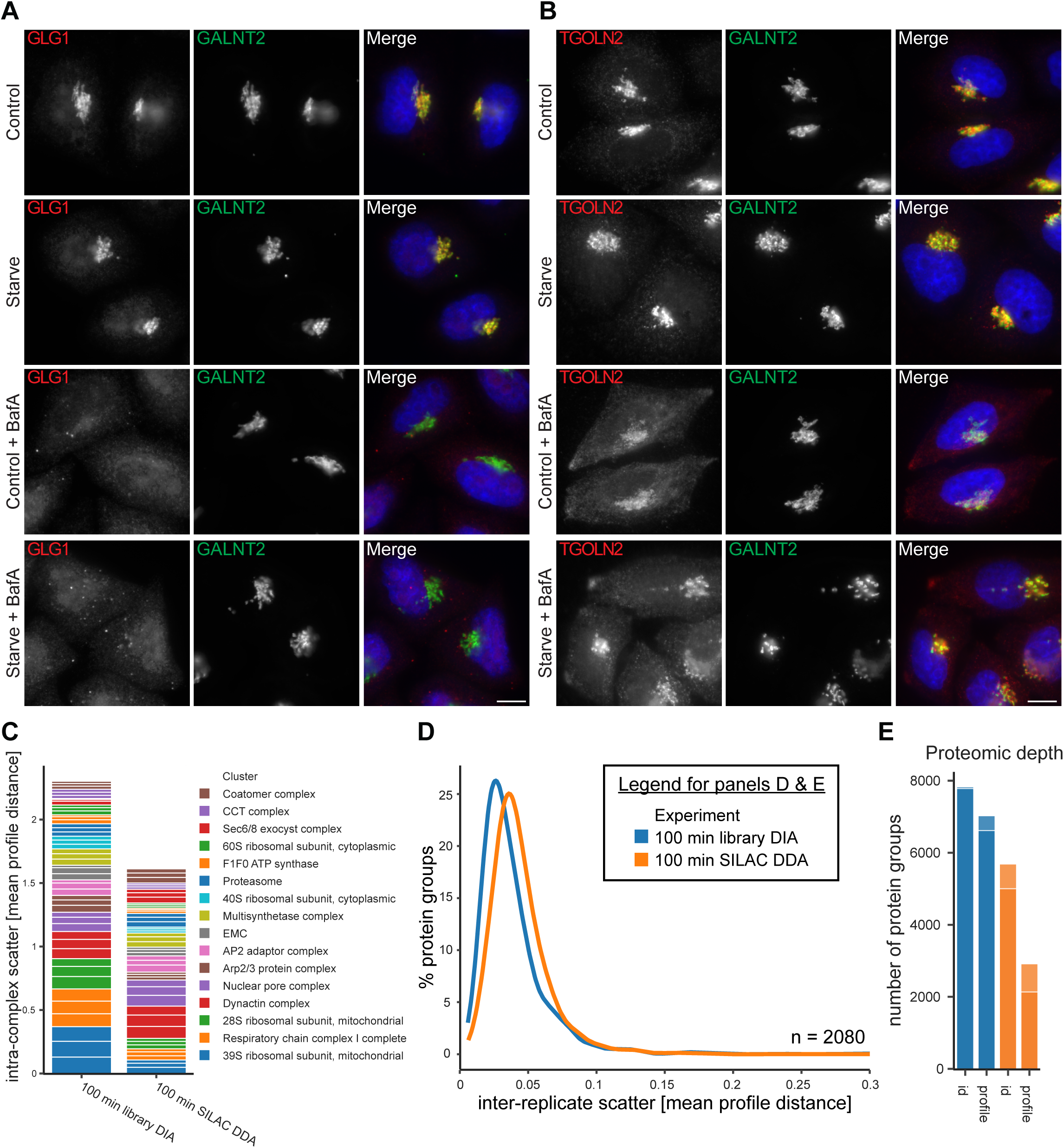
Individual effects of starvation vs. BafA treatment, and performance of DIA label-free vs. SILAC organellar maps. **A-B)** Effect of starvation and BafA treatment on Golgi protein localization. Cells were either untreated (control), starved for 1 h, treated with 100 nM BafA for 1 h (control + BafA), or starved and treated with 100 nM BafA for 1h. In the merged images, DAPI labelling of the nucleus is shown (blue). Scale bars: 10 μm. Images are representative of at least 13 images per condition. Note, the same cells are shown for the control and starve + BafA conditions as in Fig. 5G, H. **A)** Widefield imaging of immunofluorescence labelling of the Golgi proteins GLG1 (red) and GALNT2 (green). Following BafA treatment, with or without starvation, GLG1 shifts from the Golgi to endosomal structures, whereas GALNT2 remains in the Golgi. Starvation alone does not affect the localization of either protein. **B)** Widefield imaging of immunofluorescence labelling of the Golgi proteins TGOLN2 (TGN46; red) and GALNT2 (green). Following BafA treatment, with or without starvation, TGOLN2 shifts partially from the Golgi to endosomal structures, whereas GALNT2 remains in the Golgi. Starvation alone does not affect the localization of either protein. **C-E)** Performance of SILAC organellar maps. Organellar maps generated in a previous study [11,19] using SILAC metabolic labelling, with 100 min LC gradients and DDA, were evaluated against the reference single shot label-free DIA maps, with 100 min LC gradients, library processing, generated in this study (see Fig. 2). **C)** SILAC maps have lower intra-complex scatter than DIA maps, indicating higher quantification precision. **D)** DIA maps have slightly lower inter-replicate scatter than SILAC maps for the 2,080 proteins profiled across both experiment formats. **E)** DIA maps have triple the profiling depth of SILAC maps (6,615 vs 2,135 protein groups after quality filtering).

